# *Staphylococcus aureus* can use an alternative pathway to be internalized by osteoblasts in absence of β1 integrins

**DOI:** 10.1101/2023.12.13.571096

**Authors:** William Mouton, Léo-Paul Tricou, Andréa Cara, Sophie Trouillet-Assant, Daniel Bouvard, Frédéric Laurent, Alan Diot, Jérôme Josse

## Abstract

*Staphylococcus aureus* main internalization mechanism in osteoblasts relies on a tripartite interaction between bacterial fibronectin-binding proteins, extracellular matrix soluble fibronectin, and osteoblasts’ β1 integrins. Caveolins, and particularly caveolin-1, have shown to limit the plasma membrane microdomain mobility, and consequently reduce the uptake of *S. aureus* in keratinocytes. In this study, we aimed to deepen our understanding of the molecular mechanisms underlying *S. aureus* internalization in osteoblasts. Mechanistically, *S. aureus* internalization requires endosomal recycling β1 integrins as well as downstream effectors such as Src, Rac1, and PAK1. Surprisingly, in β1 integrin deficient osteoblasts, *S. aureus* internalization is restored when Caveolin-1 is absent and requires αvβ3/αvβ5 integrins as backup fibronectin receptors. Altogether, our data support that β1 integrins regulate the level of detergent-resistant membrane at the plasma membrane in a an endosomal and Caveolin-1 dependent manner.

**SUMMARY STATEMENT:** *Staphylococcus aureus* can be internalized by osteoblasts via a different mechanism than the main α5β1/fibronectin/fibronectin-binding protein that likely involves αvβ3 or αvβ5 integrin.

## INTRODUCTION

Integrins are the main class of receptors implicated in the interactions between cells and extracellular matrix (ECM) (Hynes, 1992). Binding of integrins to ECM ligands can activate signaling pathways, including the FAK, Src, Rho GTPases (Rac1, CDC42 and RhoA), Erk and PI3-kinase pathways. This interaction influences cytoskeleton organization, which accounts for many of integrin-regulated biological functions such as proliferation, apoptosis, cell fate decision, and extracellular matrix organization (Giancotti and Ruoslahti, 1999).

β1 integrins play important roles in the ECM / cell interactions, especially in bone and joint tissues. Direct interactions between osteoblasts, the bone forming cells, and the ECM, are mediated by a select group of integrin receptors that includes α5β1, α3β1 and α8β1 (Moursi et al., 1997; Brunner et al., 2013). These integrins can interact with fibronectin, a glycoprotein that can be found in the ECM, but also as a soluble form in plasma (Pankov and Yamada, 2002). It was reported that α5β1 integrin mediates critical interactions between osteoblasts and fibronectin, which are required for both bone morphogenesis and osteoblast differentiation (Moursi et al., 1997; Brunner et al., 2011).

Several studies showed that integrins are associated with detergent-resistant membrane (DRM), specific membrane microdomains that are rich in cholesterol, sphingolipids, and other proteins (Green et al., 1999; Lietha and Izard, 2020). DRM play important roles in the regulation of cell adhesion to the ECM and associated cell migration, by concentrating molecules such as Src or Rac1 involved in the downstream signaling pathways of integrins (Lietha and Izard, 2020).

Moreover, integrins, including β1 integrins, are known to be constitutively endocytosed and recycled, which are essential processes for cell migration, and to ensure the turnover of the ECM (Bretscher, 1989; Moreno-Layseca et al., 2019). While endocytosis of integrins can be mediated through clathrin-dependent pits, it has been shown that β1 integrin endocytosis requires another surface-associated unit called caveolae. These plasma membrane compartments are a subtype of DRMs rich in caveolins (Cav), a family of integral membrane proteins responsible for the caveolae-dependent endocytosis mechanism. (Shi and Sottile, 2008).

Caveolae are cholesterol-enriched domains predominantly composed of Caveolin-1 (Cav1), which is the most abundant isotype within the caveolar structure (Drab et al., 2001). Cav1 is also the key element of the caveolar endocytosis process, as it requires Cav1 phosphorylation on Tyr 14 to occur (del Pozo et al., 2005). Furthermore, Rac-binding sites can be found in cholesterol-enriched domains and seem to be endocytosed by a caveolin-dependent pathway. Consequently, Cav1 is very likely to manage regulation of Rac-binding sites endocytosis (Drab et al., 2001; Del Pozo and Schwartz, 2007).

However, this internalization is only possible in the presence of β1 integrins, as they play a major role in caveolae insertion stability into the plasma membrane (Wickström et al., 2010). On the other side of the plasma membrane, β1 integrins are also the key receptors in the integrin-mediated internalization of fibronectin-binding pathogens such as *Staphylococcus aureus* (Hoffmann et al., 2011).

*S. aureus* is a commensal bacterium of the skin that can cause a wide range of infections such as pneumonia, bacteriemia, infective endocarditis but also bone and joint infections (Lowy, 1998). For many years, *S. aureus* has been considered as an extracellular pathogen, but is now widely acknowledged to be a facultative intracellular organism. Its intracellular survival in neutrophils, monocytes, macrophages, and other professional phagocytic cells, has been first described in the 1950s (Rogers and Tompsett, 1952; Kapral and Shayegani, 1959). Intracellular *S. aureus* is protected against both antibiotics and the host immune response, using the cell as a trojan horse (Gresham et al., 2000). However, *S. aureus* internalization by? nonprofessional phagocytic cells (NPPCs) such as epithelial cells, endothelial cells, fibroblasts, or osteoblasts can also occur (Ogawa et al., 1985; Hudson et al., 1995; Bayles et al., 1998; Sinha et al., 2000). More importantly, several studies reported the presence of intracellular *S. aureus* inside NPPCs from clinical samples such as nasal mucosa samples or bone samples, suggesting that *S. aureus* presence inside host cells may contribute to its pathogenicity (Bosse et al., 2005; Clement et al., 2005).

*In vitro* experiments revealed that *S. aureus* internalization in NPPCs is mainly achieved through a tripartite interaction between bacterial adhesins called fibronectin-binding proteins (FnBP), free soluble fibronectin in the extracellular medium and α5β1 integrin on the host NPPC surface (Sinha et al., 1999; Fowler et al., 2000). *S. aureus* can express 1 or 2 FnBP isotypes (FnBPA and FnBPB), for which the deletion or an inactivation leads to a significant decrease of the internalization efficacy (Dziewanowska et al., 1999; Sinha et al., 2000). While β1 integrin downstream effectors such as Src and FAK are also involved in the NPPCs invasion pathway, it has been observed that *S. aureus* colocalizes with host membrane cholesterol, strongly suggesting that internalization happens in cholesterol-enriched microdomain such as DRMs (Agerer et al., 2003; Fowler et al., 2003; Mittal et al., 2019; Ji et al., 2020). Regarding the phagocytosis mechanism, Veiga *et al*. reported that *S. aureus* internalization in HeLa cells is clathrin-dependent (Veiga et al., 2007). Hoffman *et al*. showed that Cav1-deficient fibroblasts exhibited an increased uptake of *S. aureus*. They reported that this potentiated internalization may be related to an enhanced mobility of membrane microdomain-associated proteins when Cav1 is missing (Hoffmann et al., 2010). However, signaling pathways involved as well as roles of Cav1 and DRMs are still not fully understood.

In this study, we aim to deepen our understanding of molecular mechanisms required for *S. aureus* internalization by NPPCs. First, we validated the role of FnBPs, fibronectin and β1 integrin in our murine osteoblast model. We reported the role of Src, Rac1 and PAK1 (a downstream effector of Rac1 that regulates actin structure) in the internalization process. Then, we showed that the absence of Cav1 expression rescued the *S. aureus* internalization in β1-deficient osteoblasts. Finally, we reported that Cilengitide, an αv integrin inhibitor, significantly impaired *S. aureus* internalization in β1-and Cav1-deficient osteoblasts, suggesting that αvβ3/αvβ5 integrin may play the role of an alternative cell receptor for *S. aureus* internalization in absence of β1 integrin and Cav1.

## MATERIAL AND METHODS

### *Staphylococcus aureus* strains

We used the *S. aureus* 8325-4 and its isogenic strain *S. aureus* DU5883, which is deficient for FnBPA and FnBPB, and therefore incapable to invade cells (Greene et al., 1995; Mouton et al., 2021).

### Cell lines and culture conditions

Several murine pre-osteoblastic cell lines, obtained after immortalization of mouse primary cells with the T viral oncogene of SV40, were used in this study (**Table 1**). Cells were cultured in 75 cm^2^ flasks (T75, Corning Inc. BD Falcon, Corning, NY, USA) at 37°C under 5% CO_2_, in a complete culture medium composed of DMEM (Dulbecco’s Modified Eagle Medium) containing pyruvate, D-glucose (1g/L) and L-Glutamine (1g/L) (Thermo Fischer Scientific Gibco 31885, Waltham, MA, USA) and supplemented again with L-glutamine (Gibco, 25030), 10% fetal calf serum (FCS, Gibco 10270), and with both penicillin (100μg/mL) and streptomycin (100μg/ml) (Gibco 15140).

**Table 1:**
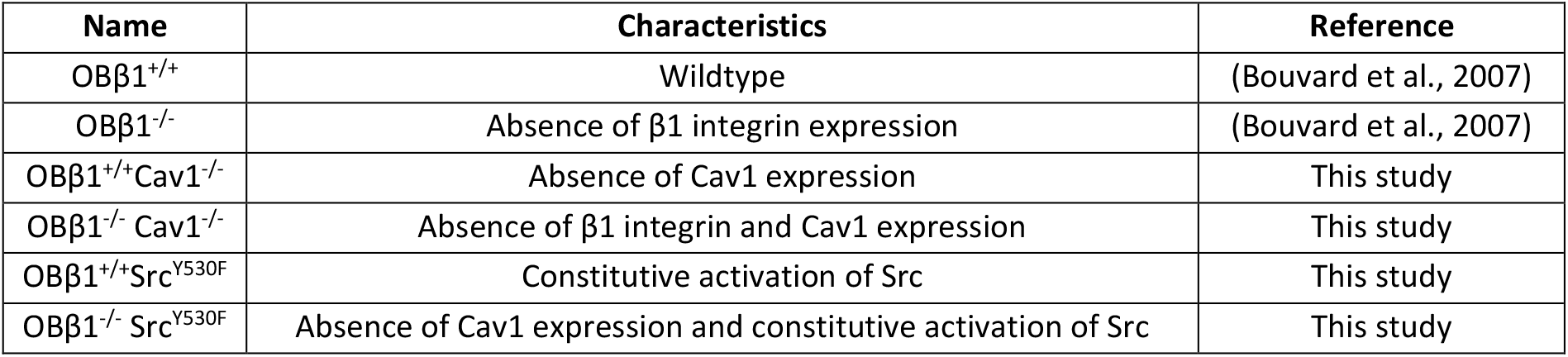
murine pre-osteoblastic cell line used in this study.

### Infection and lysostaphin protection assay

50,000 cells per well were seeded in a 24-well plate (Corning Inc. BD Falcon, Corning, NY, USA) and incubated for 48h, before being infected at a MOI of 1:1 with the previously indicated strains. After 2 hours of contact, 10µg/mL lysostaphin (Sigma-Aldrich, Burlington, MA, USA) were added for 1h to eliminate adherent and non-adherent extracellular bacteria. Bacterial internalization was then assessed after osmotic shock-induced cell lysis and plating of lysates on TSA agar plates (BioMérieux, Marcy-l’Étoile, France) using the Easy Spiral automaton. (Interscience, Roubaix, France). Colony Forming Units (CFU) were counted after 18 to 24h at 37°C.

### Antibodies and chemical inhibitors

Several chemical inhibitors were used in these experiments and are listed in **Table 2**. All inhibitors were diluted in culture medium without antibiotics. The cells were pretreated for 1h before infection and during the 2h-long contact with bacteria. Cell infection was carried out as described in the above paragraph.

**Table 2:** Chemical inhibitors used in this study.

| Inhibitor / Antibodies | Target | Used concentration (µM or µg/ml for antibodies) | Supplier |
| --- | --- | --- | --- |
| Src Inhibitor No.5 | Src | 10 | ProteinKinase (Kassel, Germany) |
| EHT 1864 | Rac1 | 5 | Sigma-Aldrich (Burlington, MA, USA) |
| p21-Activated Kinase Inhibitor III, IPA-3 | Pak1 | 5 |  |
| Primaquine | endosomal trafficking | 5 |  |
| Cilengitide | αv integrin | 0.5 |  |
| anti-CD29 antibodies | integrin β1 (CD29) | 10 | BD Pharmingen (Franklin Lakes, MA, USA) |
| anti-fibronectin antibodies | fibronectin | 10 | BD Transduction (Franklin Lakes, MA, USA) |

### Detergent Resistant Membranes distribution assessment by flow cytometry

DRMs distribution on the cell surface was assessed by staining the cells with cholera-toxin coupled to FITC (CT-FITC, Sigma-Aldrich, Burlington, MA, USA). The target of cholera toxin is GM1 (monosialotetrahexosylganglioside), a sphingolipid used as a marker of DRMs. Trypsinized cells (100,000) were washed once with ice-cold PBS (500ml-5min-4°C) and stained with 1µg/mL of CT-FITC for 30min at 4°C. The cells were then washed twice with ice-cold PBS and fixed at 4°C with 3.7% paraformaldehyde (PFA, Sigma-Aldrich, Burlington, MA, USA) for 10min. The fixed cells were then washed once in ice-cold PBS and stored in 1mL PBS at 4°C in the dark until analysis on a BD Accuri™ C6 (FL1-H).

### Graphical representation and statistical analysis

Results were presented as number of intracellular CFU for 50 000 cells, excepted for Fig. 2D and Fig. 3B where results were presented as percentage of CT-FITC staining compared to OBβ1^+/+^ cells. Data is presented as boxplots (1^st^ quartile, median, 3^rd^ quartile) with whiskers indicating minimal and maximal values. All value points are also presented for each condition. Due to the number of values, non-parametric Mann-Whitney tests were performed when applicable. All analysis were performed using GraphPad Prism 9 Software (GraphPad Software Inc., Boston, MA, USA).

## RESULTS

### Validation of the “FnBPs-fibronectin-β1 integrin” internalization pathway in OBβ1 cells

To validate the role of the β1 integrin in the cellular internalization of *S. aureus*, we used the OBβ1 murine pre-osteoblasts cell line expressing (OBβ1^+/+^) or not (OBβ1^-/-^) β1 integrin (Brunner et al., 2011) and the *S. aureus* strains expressing (8325-4) or not (DU5883) FnBPs in a lysostaphin protection assay. In absence of β1 integrin, internalization of the 8325-4 strain into osteoblasts was significantly decreased by 3 Log, from 2.2×10^3^ to 0 intracellular CFU per 50,000 cells (median values, **Fig. 1A**, p<0.0001). For the DU5883 strain, the internalization significantly dropped from 15,4 intracellular CFU per 50,000 cells to 0 in absence of β1 integrin (**Fig. 1A**, p<0.01). Regarding the FnBP-dependent internalization, the internalization of DU5883 was significantly decreased by 99% in OBβ1^+/+^ cells compared to the 8325-4 parental strain (**Fig. 1A**, p<0.0001). In OBβ1^-/-^ cells, no significant difference was observed between 8325-4 and DU5883 strains due to a negligeable number of internalized bacteria (**Fig. 1A**). The role of β1 integrin and fibronectin in the internalization process was also confirmed using anti β1 and anti-fibronectin blocking antibodies respectively **Fig. 1B, C**).

**Figure 1:**
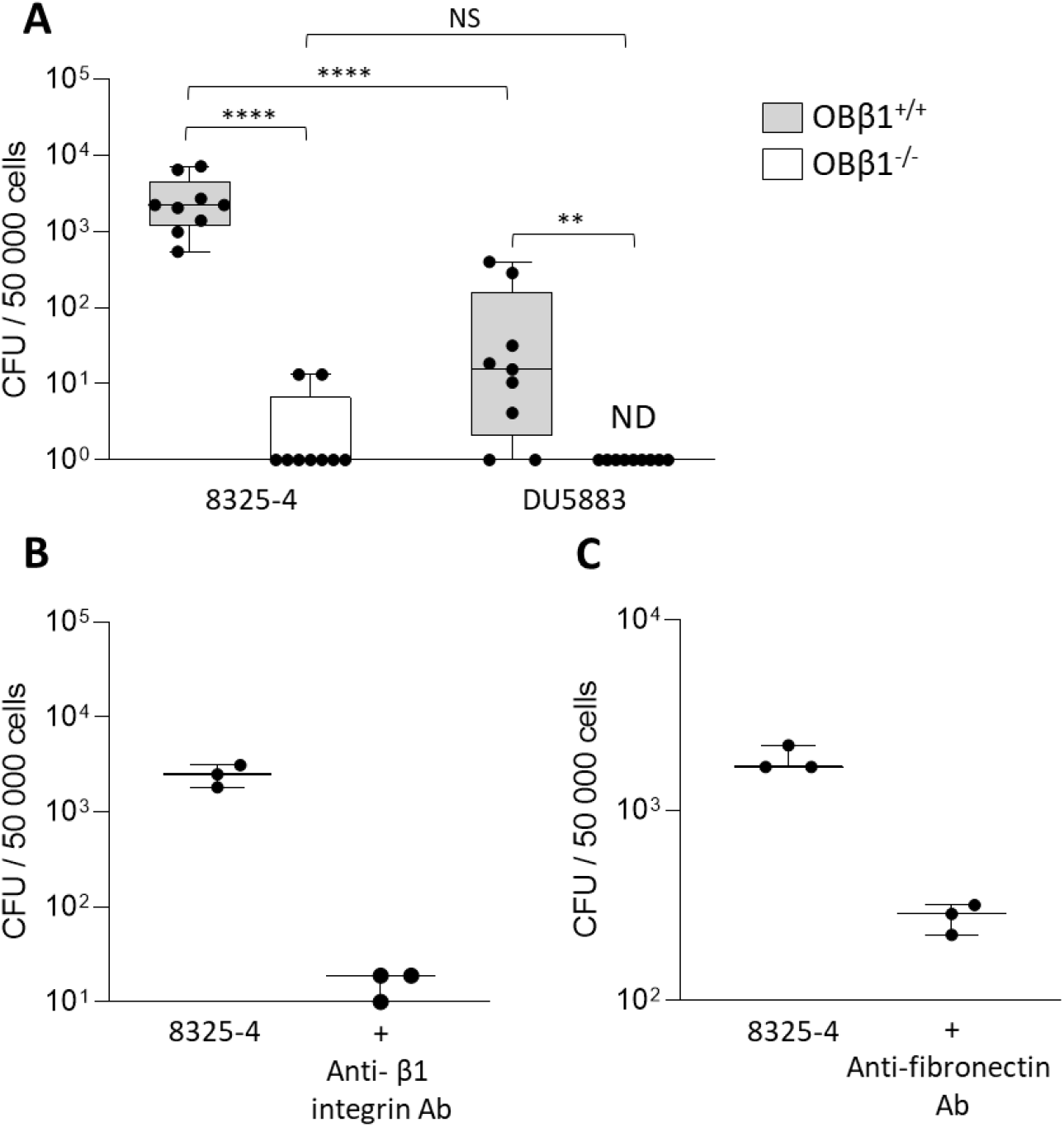
Internalization of *Staphylococcus aureus* 8325-4 inside OBβ1 osteoblasts is dependent of Fibronectin-binding proteins, fibronectin and β1 integrin. (**A**) Internalization of *S. aureus* 8325-4 and DU5883 (its isogenic counterpart deficient for FnBPs) inside OBβ1^+/+^ and OBβ1^-/-^ (OBβ1 deleted for β1 integrin). 3 independent experiments in technical triplicates (3 wells for each condition for each experiment) were performed. Non-parametric Mann-Whitney tests were performed. *, **, *** or **** means *p* < 0.05, *p* < 0.01, *p* < 0.001 and *p* < 0.0001, respectively. (**B**) Internalization of *S. aureus* 8325-4 inside OBβ1^+/+^ pretreated or not with anti-β1 integrin antibodies before the infection. (**C**) Internalization of *S. aureus* 8325-4 inside OBβ1^+/+^ pretreated or not with anti-fibronectin antibodies before the infection. Results were presented as number of intracellular colony-forming units (CFU) for 50 000 cells. As validation experiments, only one experiment in technical triplicate was performed for (**B**) and (**C**), so no statistical analysis was performed.

### Src, Rac1, PAK1, and endosomal recycling are involved in the internalization of *S. aureus* 8325-4 in OBβ1 cells

The signaling protein Src has been shown to play a pivotal role in the internalization process of *S. aureus* in HeLa and 293T cells (Agerer et al., 2003; Fowler et al., 2003). When OBβ1 cells were treated with a Src inhibitor, the internalization of 8325-4 was strongly impaired, endorsing that Src activity is important for the internalization of *S. aureus* (2.6×10^3^ vs. 1.5×10^2^ intracellular CFU per 50,000 cells, median values, p<0.0001, **Fig. 2A**). As former studies have suggested that Src activity was triggered following the contact between *S. aureus* and β1 integrin (Fowler et al., 2003), we also assessed if Src activation could rescue the internalization of *S. aureus* in absence of β1 integrin, by expressing a constitutively active form of Src (Src^Y530F^). We observed that constitutive activation of Src decreased the internalization of 8325-4 strain, from 2.5 ×10^3^ to 5.8×10^2^ intracellular CFU for 50,000 cells (**Fig. S1**, median values, p<0.0411). In OBβ1^-/-^Src^Y530F^, constitutive activation of Src was not able to rescue *S. aureus* internalization in absence of β1 integrin (**Fig. S1**).

**Figure 2:**
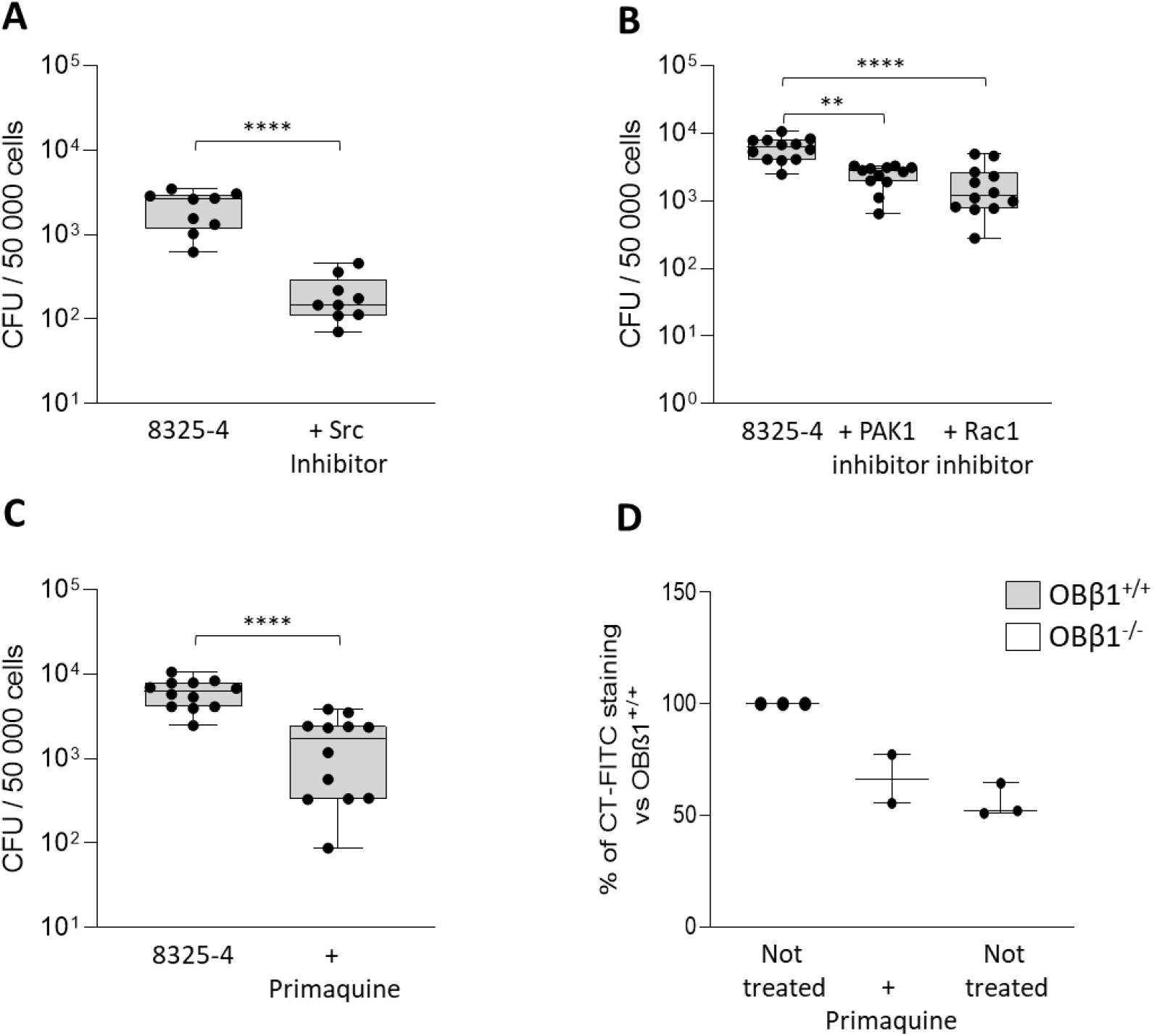
Internalization of *Staphylococcus aureus* 8325-4 inside OBβ1 osteoblasts is dependent of Src kinase, Rac1, PAK1 and endosomal recycling. (**A**) Internalization of *S. aureus* 8325-4 inside OBβ1^+/+^ pretreated or not with Src inhibitor before the infection. (**B**) Internalization of *S. aureus* 8325-4 inside OBβ1^+/+^ pretreated or not with Rac1 and PAK1 inhibitors prior to the infection. (**C**) Internalization of *S. aureus* 8325-4 inside OBβ1^+/+^ pretreated or not with primaquine before the infection. Results were presented as number of intracellular colony-forming units (CFU) for 50 000 cells. At least 3 independent experiments in technical triplicates (3 wells for each condition for each experiment) were performed. Non-parametric Mann-Whitney tests were performed. *, **, *** or **** means *p* < 0.05, *p* < 0.01, *p* < 0.001 and *p* < 0.0001, respectively. (**D**) Presence of GM1 (marker of DRMs) at the surface of OBβ1^+/+^, OBβ1^+/+^ pretreated with primaquine and OBβ1^-/-^. Results were presented as percentage of CT-FITC staining compared to the non-treated OBβ1^+/+^ cells. Three independent experiments but without technical duplicate or triplicate were performed for (**D**), so no statistical analysis was performed.

Then we tested the role of Rac1 and PAK1 regarding the internalization of *S. aureus*. These two proteins are downstream Src and are involved in several cell mechanisms, especially in the cytoskeletal reorganization. Using chemical inhibitors, we observed that inhibition of PAK1 reduced *S. aureus* internalization by 56% (6.3×10^3^ vs. 2.8×10^3^ intracellular CFU per 50,000 cells, median values, in presence or absence of PAK1 inhibitor, respectively, p<0.01, **Fig. 2B**). In the same way, Rac1 inhibition also impaired *S. aureus* internalization, decreasing *S. aureus* internalization by 81% (6.3×10^3^ vs. 1.2×10^3^ intracellular CFU per 50,000 cells, median values, p<0.0001, in presence or absence of Rac1 inhibitor, respectively, **Fig. 2B**).

Finally, we investigated the role of DRMs and endosomal recycling on the internalization of *S. aureus*, as β1 integrin and DRMs co-regulate their recycling at the plasma membrane (Leitinger and Hogg, 2002). After its endocytosis, integrin and DRMs can be recycled to the membrane. To do so, OBβ1^+/+^ cells were pretreated with primaquine, an anti-malaria drug that interferes with membrane recycling from endosomes to the plasma membrane through a direct interaction with them (van Weert et al., 2000) After primaquine treatment, *S. aureus* 8325-4 strain internalization was decreased by 72% (6.3×10^3^ vs. 1.8×10^3^ intracellular CFU per 50,000 cells, median values, p<0.0001, **Fig. 2C**). To support the impact of primaquine on endosomal recycling, we used our ability to stain DRMs (especially glycosphingolipid GM1) using cholera-toxin fused to FITC fluorescent protein (CT-FITC) to assess the integrity of the vesicular transport in our OBβ1^+/+^ and OBβ1^-/-^ cell lines. Our results showed that primaquine treatment induced a 34% decreased of DRMs presence at the cell surface compared to non-treated cells (**Fig. 2D**). Interestingly, we also observed that β1 integrin deletion led to a 48% decrease of CT-FITC staining compared to the non-treated wildtype cells (**Fig. 2D**).

Overall, inhibiting the downstream effectors of integrin β1 or its putative relocation to cell membrane through endosomal recycling impacts the internalization of *S. aureus*. Moreover, it seems that β1 integrin might be involved in the control of the recycling endosome trafficking back to the plasma membrane. Altogether, these observations suggest that inhibition of β1 integrin or its supposed relocation to the plasma membrane via endosomal recycling appears to impair *S. aureus* internalization.

**Figure S1:**
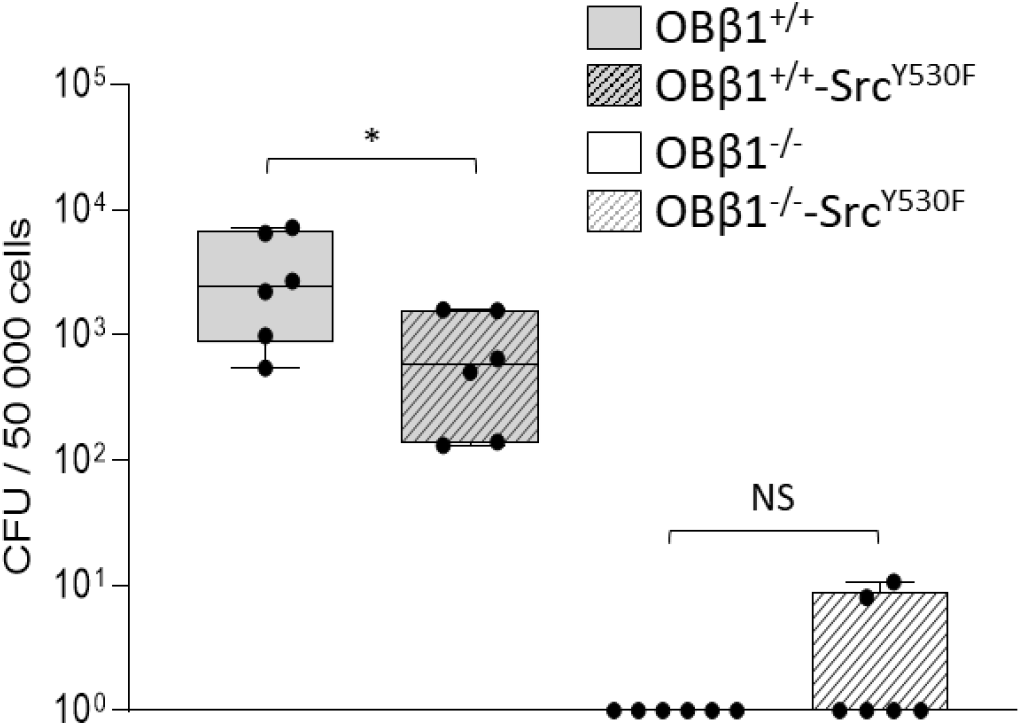
Constitutive activation of Src was not able to rescue the internalization of *Staphylococcus aureus* 8325-4 in OBβ1^-/-^ cells. Internalization experiments were performed using OBβ1^+/+^, OBβ1^+/+^Src^Y530F^ (with a constitutive activation of Src), OBβ1^-/-^ and OBβ1^-/-^Src^Y530F^. Results were presented as number of intracellular colony-forming units (CFU) for 50 000 cells. Two independent experiments in technical triplicates (3 wells for each condition for each experiment) were performed. Non-parametric Mann-Whitney tests were performed. *, **, *** or **** means *p* < 0.05, *p* < 0.01, *p* < 0.001 and *p* < 0.0001, respectively.

### Caveolin-1 deletion partially rescues the internalization in OBβ1^-/-^ cells

Caveolin-1 (Cav1) is a structural component of membrane compartments called caveolae that has been thought to be a negative regulator of the internalization pathway in fibroblasts expressing β1 integrin (Hoffmann et al., 2010). We decided to analyze how Cav1 modulates *S. aureus* internalization in OBβ1 cells, especially in absence of β1 integrin. Contrary to observations in fibroblasts, we did not observe an increased bacterial internalization when the caveolin gene was invalidated in OBβ1^+/+^ cells (**Fig. S2**). Interestingly, while we observed 0 CFU for OBβ1^-/-^ cells with functional Cav1, we noticed a significant rescue of *S. aureus* internalization with 3,4×10^2^ intracellular CFU per 50,000 cells in Cav1-deficient OBβ1^-/-^, which accounts for 12% of *S. aureus* internalization in OBβ1^+/+^ cells (3.4×10^2^ vs. 2.8×10^3^ intracellular CFU per 50,000 cells, median values, **Fig. 3A**). GM1 and DRMs presence among this these 3 cell types were then assessed using CT-FITC staining. A rescue of DRMs presence at the host cell surface was also observed as OBβ1^-/-^ Cav1^-/-^ cells presented a 115% of CT-FITC staining similar to OBβ1+/+ cells, whereas staining in OBβ1^-/-^ cells was decreased by 48% (median values, **Fig. 3B**).

**Figure 3:**
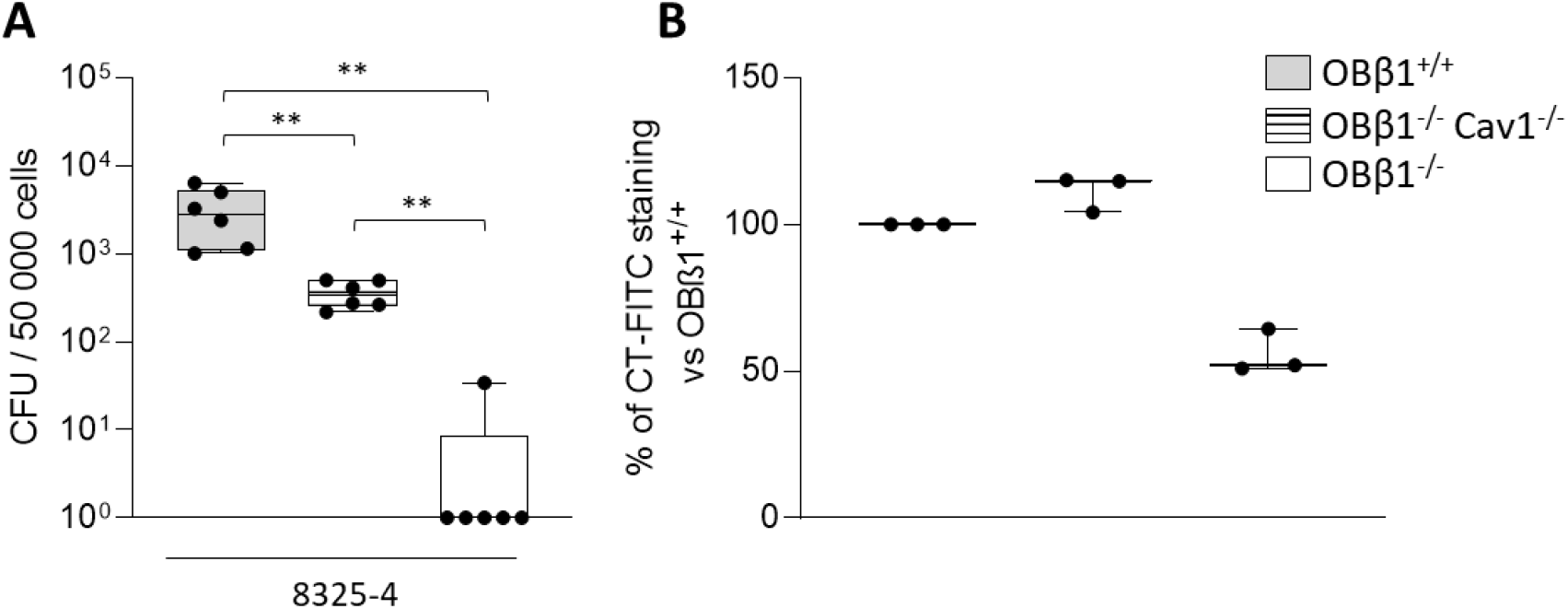
Absence of Cav1 in OBβ1^-/-^ (OBβ1^-/-^Cav^-/-^ osteoblasts) rescues the internalization of *Staphylococcus aureus* 8325-4. (**A**) Internalization of *S. aureus* 8325-4 inside OBβ1^+/+^, OBβ1^-/-^Cav1^-/-^ and OBβ1^-/-^. Results were presented as number of intracellular colony-forming units (CFU) for 50 000 cells. Two independent experiments in technical triplicates (3 wells for each condition for each experiment) were performed. Non-parametric Mann-Whitney tests were performed. ** means *p* < 0.01. (**B**) Presence of GM1 (marker of DRMs) at the surface of OBβ1^+/+^, OBβ1^-/-^ Cav1^-/-^ and OBβ1^-/-^. For panel B, results were presented as percentage of CT-FITC staining compared to the control cells (OBβ1^+/+^). Three experiments but without technical duplicate or triplicate were performed, thus no statistical analysis was performed.

Our results suggest that *S. aureus* could be internalized via an alternative pathway in absence of β1 integrin, which is potentially repressed by Cav1. Interestingly, Cav1 also impacts the endosomal recycling in OBβ1^-/-^ cells.

**Figure S2:**
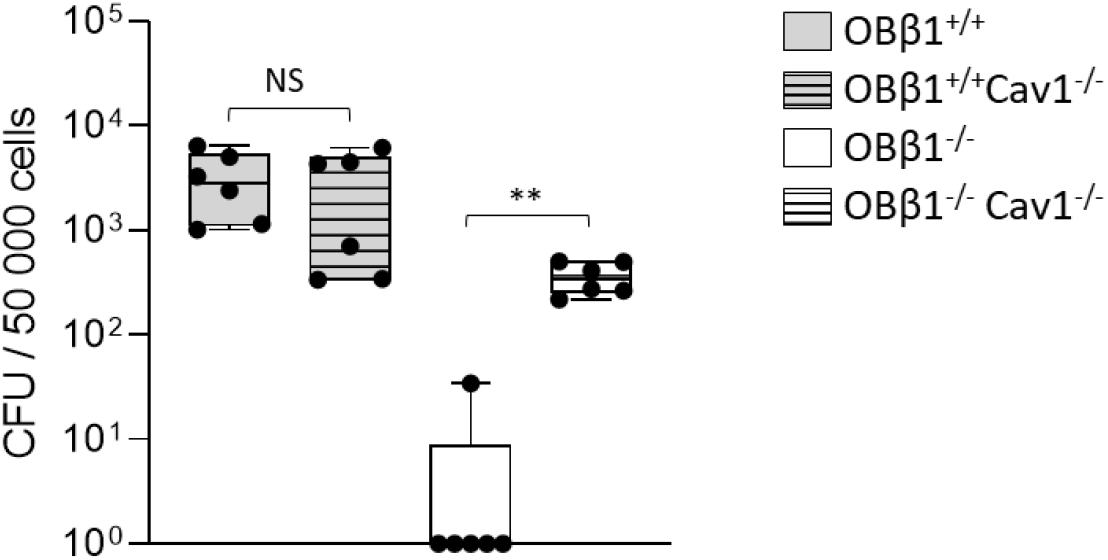
Caveolin1 deletion in OBβ1^+/+^ did not impact the internalization of *Staphylococcus aureus* 8325-4. Internalization experiments were performed using OBβ1^+/+^, OBβ1^+/+^Cav1^-/-^, OBβ1^-/-^ and OBβ1^-/-^Cav1^-/-^. Results were presented as number of intracellular colony-forming units (CFU) for 50 000 cells. At least 2 independent experiments in technical triplicates (3 wells for each condition for each experiment) were performed. Non-parametric Mann-Whitney tests were performed. *, **, *** or **** means *p* < 0.05, *p* < 0.01, *p* < 0.001 and *p* < 0.0001, respectively.

### Internalization of *S. aureus* in absence of β1 integrin and Cav1 is FnBP-dependent, αvβ3-dependent, and activates a similar intracellular signalization as in β1 integrin-dependent internalization

To characterize the internalization pathway in OBβ1^-/-^Cav1^-/-^ cells, we first performed internalization experiments using *S. aureus* strains with (8325-4) or without (DU5883) functional FnBPs. We observed that the internalization of DU5883 is decreased by 98% compared to 8325-4 in OBβ1^-/-^Cav1^-/-^ cells (5.5×10^2^ vs. 13 intracellular CFU per 50,000 cells, median values, p<0.0001, **Fig. 4A**). This pattern is similar to the one observed for OBβ1^+/+^ cells, suggesting that FnBPs are involved (7.6×10^3^ vs. 83 intracellular CFU per 50,000 cells, median values, p<0.0001, **Fig. 4A**). Then, we identified the host cell receptor that mimics the role of the α5β1 integrin when β1 integrin is missing. As the αvβ3 integrin was reported in *S. epidermidis* adhesion on osteoblasts, we hypothesized that it could be an adequate host cell receptor (Claro et al., 2015). For this purpose, we utilized Cilengitide, a specific inhibitor of αvβ3 and αvβ5 integrins (Hariharan et al., 2007). After treatment of OBβ1^-/-^Cav1^-/-^ cells with Cilengitide, the internalization of *S. aureus* 8325-5 was decreased by 90% (1.38×10^2^ vs. 13 intracellular CFU per 50,000 cells, median values, p<0.01, **Fig. 4B**). However, when OBβ1^+/+^ cells were treated with Cilengitide, no effect on the internalization of *S. aureus* was observed (**Fig. 4B**). Considering these results, it seems that αvβ3 integrin may be the major host receptor involved in internalization of *S. aureus* in OBβ1 cells, in absence of β1 integrin and Cav1. Finally, we tested the role of host proteins involved in the intracellular signalization and cytoskeleton reorganization. Thanks to chemical inhibitors, we observed that Src, Rac1, PAK1, and the endosomal recycling (through primaquine treatment) were also involved in the internalization of *S. aureus* in OBβ1^-/-^Cav1^-/-^ cells, as observed previously for OBβ1^+/+^ cells (**Fig. 4C**). In OBβ1^-/-^Cav1^-/-^ cells, the internalization of *S. aureus* was significantly decreased after Src inhibition, decreased by 73% after PAK1 inhibition, 60% after Rac1 inhibition, and 82% after primaquine treatment (**Fig. 4C**).

**Figure 4:**
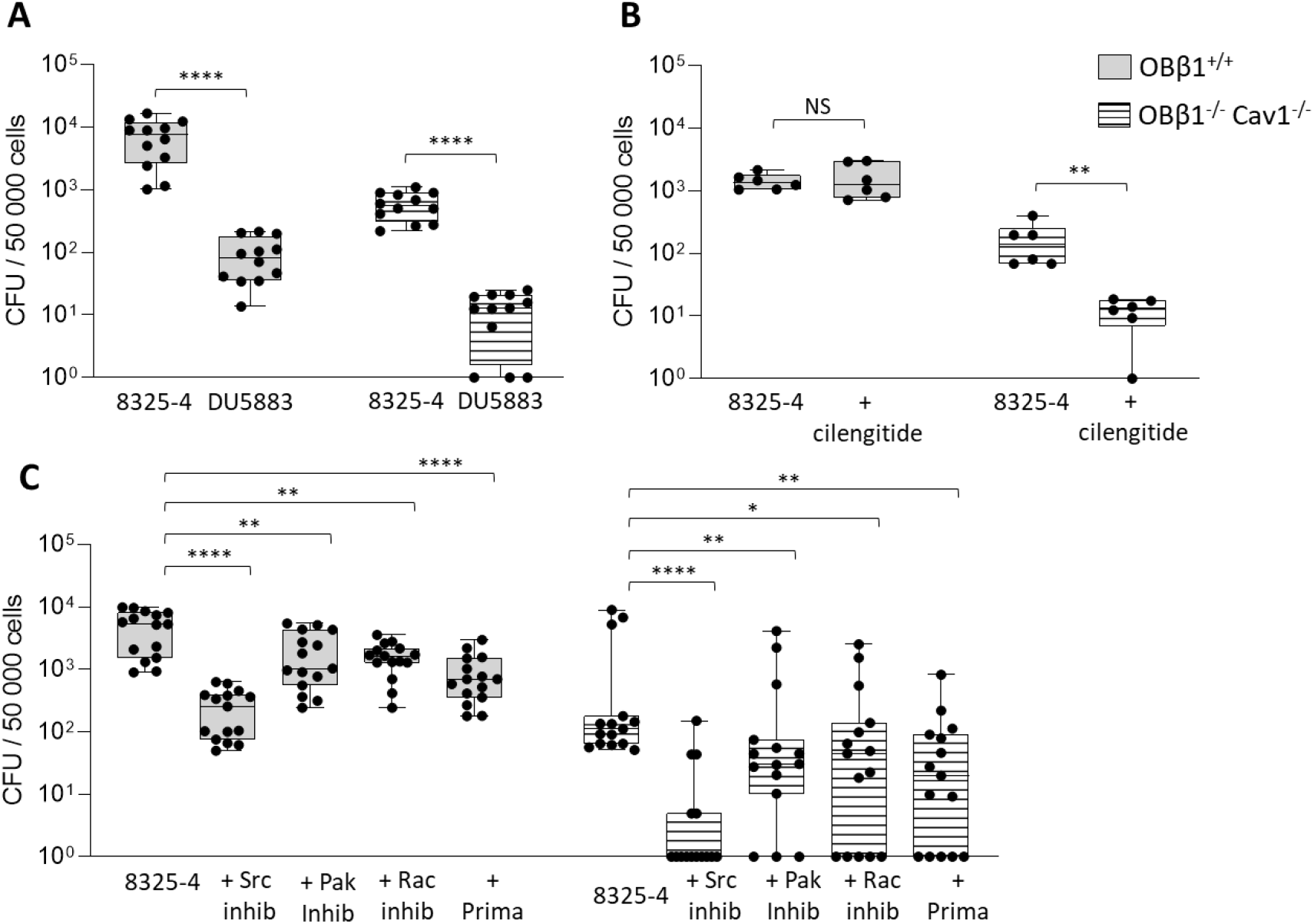
Internalization of *Staphylococcus aureus* 8325-4 inside OBβ1^-/-^Cav1^-/-^ cells is dependent on Fibronectin-binding proteins, αvβ3 integrin, Src kinase, Rac1, PAK1 and endosomal recycling. (**A**) Internalization of *S. aureus* 8325-4 and DU5883 inside OBβ1^+/+^ and OBβ1^-/-^Cav1^-/-^. (**B**) Internalization of *S. aureus* 8325-4 inside OBβ1^+/+^ and OBβ1^-/-^Cav1^-/-^ pretreated or not with Cilengitide (a specific inhibitor of αv integrins) before the infection. (**C**) Internalization of *S. aureus* 8325-4 inside OBβ1^+/+^ and OBβ1^-/-^ Cav1^-/-^ pretreated or not with Src, PAK1 and Rac1 inhibitors or primaquine before the infection. Results were presented as number of intracellular colony-forming units (CFU) for 50 000 cells. At least 2 independent experiments in technical triplicates (3 wells for each condition for each experiment) were performed. Non-parametric Mann-Whitney tests were performed with Prism GraphPad. *, **, *** or **** means *p* < 0.05, *p* < 0.01, *p* < 0.001 and *p* < 0.0001, respectively.

## DISCUSSION

*S. aureus* infection often leads to difficult to treat bone and joint infections, mainly because of its ability to escape the immune response via internalization into osteoblasts. This internalization not only perturbs bone deposition balance by reducing osteoblast differentiation, and promoting bone degradation, but also constitutes a bacteria reservoir promoting chronicity (Clement et al., 2005; Tuchscherr et al., 2010; Mouton et al., 2021). In this study, we aimed to deepen our current understanding of the molecular mechanisms that control *S. aureus* internalization (Josse et al., 2015). Although we confirmed previous data showing that *S. aureus* internalization into osteoblasts requires a functional β1 integrin/fibronectin/FnBP tripartite axis, our present work provides a novel and extended view on how β1 integrins are involved in this process. Indeed, we propose a new picture in which β1 integrins promote *S. aureus* internalization not only as cell surface receptor but also by driving GM1 relocation to the plasma membrane. GM1, is a lipid marker for the so-called detergent resistant rich membranes (DRM), involved in the formation of membranous subcellular domains that concentrate and stabilize signaling proteins such as Rac1, PAK1, and Src. Consequent to this local recruitment and activation of signaling proteins, numerous biological processes are triggered such as cell adhesion, migration, and internalization (Josse et al., 2017).

Our data fully supports previous data showing that β1 integrins are required for GM1 trafficking back to the plasma membrane (Wickström et al., 2010). Loss of cell adhesion to the ECM has shown to promote DRM internalization likely via a Cav1 dependent route. Indeed, the low GM1 content associated with the absence of β1 integrins was partially restored back upon co-removal of Cav1. Thus, our data support this hypothesis, and demonstrate that β1 integrins, by promoting the trafficking back to and likely the stabilization of Cav1 at the plasma membrane are a key regulator of DRM dynamics at the plasma membrane. Importantly, pathophysiological consequences of β1 integrins-dependent DRM stabilization at the plasma membrane have been poorly addressed so far. Here, using an established osteoblastic model for *S. aureus* internalization, we extended these findings with a more clinical perspective. Of importance, we also discovered the role of the αvβ3 integrin as a “backup” cell receptor for *S. aureus* invasion in β1-integrin and Cav1-deficient osteoblasts, suggesting the existence of another invasion mechanism (**Fig. 4B**). It is well recognized that *S. aureus* main internalization pathway is dependent on β1 integrins via its interaction with fibronectin. Moreover, when β1 integrins are deleted, it cannot be compensated by other fibronectin integrin receptors such as αvβ3 and/or αvβ5, although being expressed by osteoblasts. Surprisingly, deletion of both β1 integrins and Cav1 restored bacterial internalization, challenging the current concept. Indeed, one of the main observations from this work was to identify β1 integrins being required for DRM localization/stabilization at the plasma membrane as an essential regulator of *S. aureus* cell invasion.

The *in vivo* interaction between *S. aureus* and bone tissue via the α-hemolysin (Hla) *S. aureus* virulence factor induced an increased Cav1 expression and a Cav1-induced DRM recruitment at the plasma membrane (Liu et al., 2020). It was also described that a decreased Cav1 expression reduced DRM concentrations at the plasma membrane, which may, based on the observations from the literature, have a negative impact on *S. aureus* internalization (Grosse et al., 1998; Lee et al., 2016). However, our results suggest the opposite, where the absence of Cav1 rescued or did not decrease *S. aureus* internalization. Corroborating our results and Hoffman’s observations in fibroblasts, Hla overexpression, which is responsible for higher Cav1 and DRM concentrations at the plasma membrane, decreased fibronectin-mediated *S. aureus* internalization by A549 lung epithelial cells (Liang and Ji, 2007; Hoffmann et al., 2010). As DRMs are cholesterol-enriched membrane structures, Hoffmann’s theory stating that the absence of Cav1 can increase mobility of membrane microdomain-associated proteins and facilitate *S. aureus* internalization seems to be more in accordance with our results (Hoffmann et al., 2010). It is however important to note that our study is restricted to an *in vitro* osteoblast internalization model. The absence of other bone tissue components like osteoclasts, osteocytes, or the extracellular mineralized matrix prevents us to conclude with certainty that our observations will remain the same *in vivo*. Thus, it is possible that *S. aureus* interactions with bone tissue *in vivo* and in the *in vitro* model of this study are different. Interestingly, the discrepancies observed between our results and the literature mainly concerns *in vivo* studies, where opposite conclusions regarding the implication of Cav1 and DRM in the internalization process of *S. aureus* have been made. On the other hand, observations by Hoffman *et al*. or Liang and Ji that are in accordance with our results were made using *in vitro* model (Liang and Ji, 2007; Hoffmann et al., 2010). Also, the genetically modified osteoblastic cell lines used in our study do not represent the cell population existing within living bone tissue. Consequently, this study only gives a first insight into the existence of an accessory *S. aureus* internalization mechanism *in vitro*. This suggest that the existence of this alternative αvβ3/5-mediated internalization pathway in *vivo* needs to be investigated to determine its relevance for potential therapeutic target.

A few other studies have tried to better decipher the internalization mechanisms by identification of accessory receptors like the major staphylococcal autolysin (Atl) and the heat shock cognate protein (Hsc70) or the human heat shock protein Hsp90 as host cell receptor (Hirschhausen et al., 2010; Tribelli et al., 2020). Regarding Hsc70, it colocalizes with fibronectin and promotes *S. aureus* internalization, suggesting that this latter acts as a bridge between Hcs70 and α5β1 integrin (Schlesier et al., 2020). However, this pathway appears more likely to be an accessory mechanism to the canonical α5β1-dependent route described here. Eventually, Jang *et al*. also observed *S. aureus* internalization in HaCaT cells via the bacterial ESAT-6-like protein EsxB (Jang et al., 2021).

Although these studies do not have a direct clinical impact to prevent or treat *S. aureus* infections, they pave the way to a new non-antibiotic approach to *S. aureus* infection treatment, by highlighting new host cell therapeutic targets. Indeed, because of *S. aureus* high propension to become resistant to current antimicrobial strategies, it would be relevant to directly focus on host cell targets instead of the bacterium (Evans and McDowell, 2021). Also, by exploring drug repurposing, time and money could be saved by using already available molecules that can act on newly discovered therapeutic targets. This could be an easy-to-implement, faster and safer alternative to new antimicrobial development in the threatening context of *S. aureus* antibiotic multiresistance (Tacconelli et al., 2018). This approach has already been reviewed by Evans and McDowell, where statins were used as prophylactic agents and adjuvant therapy against intracellular *S. aureus* in different clinical studies (Evans and McDowell, 2021). Statins have shown to reduce the risk of contracting *S. aureus* infections and improve survival rate to infections by disturbing the integrity of the host cell membrane. The comprehension of statins’ mechanism of action in the internalization pathway led to the development of a more specific molecule with similar performance and less adverse effects than statins in the context of *S. aureus* infections.

In summary, in absence of β1 integrins, but also upon recovery of DRMs at the plasma membrane, *S. aureus* internalization can be mediated by alternative fibronectin receptors such as αvβ3/5, showing that β1 integrin is not an absolute receptor for *S. aureus*. Of course, we do not underestimate the functional role for β1 integrins as an important cell surface receptor, but our work with the use of genetically modified cells (deficient in β1 integrins, Cav1 and/or both) has highlighted a double role of β1 integrins in *S. aureus* internalization process. These new findings pave the way for alternative therapeutic strategy targeting the recycling of DRMs rather than β1 integrins that are key molecules involved in many important cell functions.

## ACKNOWLEDMENTS

We acknowledged Allison Faure for her technical assistance.

## DATA AVAILABILITY

All relevant data can be found within the article and its supplementary information.

